# *Brassica napus* L. dwarfing gene: Determining candidate intervals of dwarfing genes by BSA and SNP typing

**DOI:** 10.1101/2020.08.19.256958

**Authors:** Luo Jing, Li Chao, Zhang Ruimao, Chen Zhineng, Zhang Xianqiang, Gao Zhihong, Lei Lei, Li Pan

## Abstract

The plant height of rapeseed is one of the important factors that affects the production of rapeseed. If the plant height of rapeseed is too high, on the one hand, it will cause rapeseed to lodge and affect the yield, on the other hand, it will also affect the mechanized harvesting of rapeseed. In this research, the high-stalked line (YY50) and the dwarfed line (DW871) are crossed to obtain an F2 rapeseed population which was used to build pools, and then we used this to mine the main dwarfing genes. In the pools composed of tall and short stalks, we obtained 192.80Mb clean reads, which can be used for BSA (bulked segregant analysis). Preliminary positioning around the candidate section identified 23 SNP markers. Then 17 polymorphic SNP markers were obtained through polymorphism screening. Further we narrowed the candidate interval, and finally determined between 15.51-16.60Mb of ChrA10. Through identifying 231 genes from the above interval, it’s predicted that the production of dwarf traits may be related to lignin synthesis and limited inflorescence. It provides a basis for further mapping and cloning of the dwarfing gene DW871.

## 1. Introduction

Rapeseed is an important oilseed economic crop. As China’s main imported commodity, it is also the main exporting commodity of economies such as Canada and the European Union. In 2017/2018, the world’s rapeseed production reached 74.92 MT^[1]^. The main rapeseed producing areas are Canada, the European Union, and China. The planting area in 2017/2018 was 9.5 million hm^2^, 6.6 million hm^2^, and 6.8 million hm^2^. By 2021, the global rapeseed harvest area is expected to reach 35.14 million hm^2[2]^. The research of crop dwarf breeding is one of the main methods of the green revolution. Rapeseed is another crop that dominates the green revolution besides rice and wheat^[3]^. By breeders’ long-term efforts, production of rapeseed (*Brassica napus* L.) has been improved significantly. Correspondingly, lodging restricts the increase in yield and becomes one of the main factors in the reduction of rapeseed production. Therefore, it becomes a new goal for breeders that breeding dwarf varieties that are resistant to lodging and adapt to mechanized harvesting.

The earliest discovery and report of the dwarf mutants of rapeseed found by Williams and Hill^[4]^, named Bn5-2 and Bn5-8. In China, Shi^[5]^ gets dwarf mutant ds-1 and ds-2 by EMS (Ethylmethylsulfon, 0.25%) treated breeding material. Zhang^[6]^ obtained the dwarf line DW871 through cross configuration breeding, which can be inherited stably. DW871 is a semi-dwarf variety which has definite inflorescence. According to different planting methods, the average plant height is between 70cm-139.1cm. It’s more total leaf numbers, shorter internodes, thicker xylem, lower branch position, and more compact plant type compare to wild type control. By identifying the dwarfing gene of DW871, we can make better use of this excellent germplasm resource and provide help for subsequent breeding work.

Identifying molecular markers and possible candidate genes related to the dwarfing traits of rapeseed is an important means for us to make full use of germplasm resources to increase yield and mechanization rate^[7]^. When traits are controlled by a single gene, molecular marker location is our main means to find specific traits and assisted breeding. QTL (quantitative trait locus) is often used in association analysis related to traits. By associating the marker with the target trait, we can analyze the degree of correlation between the relevant data. Before we proceed to the next step, such as gene mapping, functional analysis and gene cloning, we need to find molecular markers that are closely related to the target traits^[8]^.

At present, many results have been achieved in the research of dwarfing genes in rapeseed. These dwarfing genes affect plant height features of rapeseed through many ways such as hormone regulation, polyamines, plant cell wall synthesis, and transcriptional regulation^[9–16]^. By QTL mapping, Liu^[5]^ mapped the dwarfing gene *ds-1* between SSR markers SR12156 and SCM07, with a distance of 2.8cm from SR12156 and 0.7cm from SCM07. Wang^[17]^ used the dwarfing material M176 and found the dwarfing gene located between SSR markers Na10G06 and BrA09Y21 on the ninth chromosome. The distance from Na10G06 is 7cm and the distance from BrA09Y21 is 6.5cm. Liao^[18]^ studied the mutant GA2ox6 and determined that the gene was located on *BnaA10g1070D* on chromosome A10.

The second-generation sequencing technology (next-generation sequencing, NGS) is based on the enzymatic cascade chemiluminescence reaction of four enzymes (DNA polymerase, sulfurylase, luciferase, adenosine triphosphatase) in the same PCR reaction system, called pyrosequencing too^[19]^. BSA (Bulked Segregant Analysis) based on the hybridization of parents with significant differences in traits. In the segregating population of offspring, a certain amount of phenotypic differences is selected and DNA is mixed to construct two pools. Because the selected object has only the target trait, only the region containing the target gene at the genome level is different between the two pools, and this genetic difference can be used as a molecular marker linked to the target trait^[20]^. The emergence of NGS and BSA led to the development of NGS-based BSA analysis methods, such as QTL-seq, Mutmap, SHOREmap, etc. They are widely used in gene location and cloning^[21–23]^. By QTL-seq to perform high-depth sequencing of the DNA pool of extreme traits, and then by analyzing the Δ(SNP-index) and G’ value, we can determine the candidate interval of the DW871 dwarf gene^[24,25]^.

SNP (single nucleotide polymorphism) is a new type of molecular marker widely used in character identification of rapeseed. Most SNPs are biallelic, genetically stable, and have high reproducibility^[26]^. Compared with other types of molecular markers such as SSR, polymorphism information contained in SNPs is lower, but application of high-throughput sequencing technology makes it possible to identify a large number of SNPs in multiple genomes at the same time. And provide conditions for the development of markers for identification traits, genetic diversity and mapping^[27]^. In addition, the emergence of SNP-related technologies such as SNP chips and SNP arrays that can be used for genotyping have expanded the application range of SNP molecular markers^[28,29]^. Positional cloning is also called map-based cloning. It is a method of separating genes based on the strategy of forward genetic, based on the genetic and physical map of the phenotype of the mutant individual and the phenotype site^[30]^. For example, Malmberg^[31]^ used low-coverage nanopore long-read sequencing technology to evaluate the SNP genotyping of DH(doubled haploid) rapeseed. Sun^[32]^ used SNP molecular markers to analyze QTLs related to the weight and shape of rapeseed seeds.

DW871 is a new dwarf rapeseed strain with excellent shape. By backcrossing and crossing with other *Brassica napus* high-stalk materials to calculate the segregation ratio of progeny, we found that the plant height trait of DW871 is a dominant pair gene control, there is no cytoplasmic effect, and it is affected by modified genes, which is manifested as the genetic characteristics of quantitative traits^[33]^. In this research, QTL-seq and BSA were used to analyze DW871 to determine the candidate interval of its dwarf trait^[34]^.

## 2. Material and method

### 2.1 plant material

In this experiment, the dwarf *Brassica napus* L. DW871 was used as the experimental material and named 312bulk1, and the high-stalk material YY50 was named 312bulk2 as the control material(Fig 1). The environment has a great influence on the plant height traits of *Brassica napus* L., and the materials used are all taken from the Guizhou Academy of Agricultural Sciences which lives in Huaxi District, Guiyang, Guizhou Province, 106°59’ east longitude and 26°01’ north latitude. This place belongs to a subtropical humid and temperate climate, with an altitude of 1150m, an average precipitation of 84.4mm, an average temperature of 13.70°C, and a sunshine duration of 100.90h.

**Fig 1.**
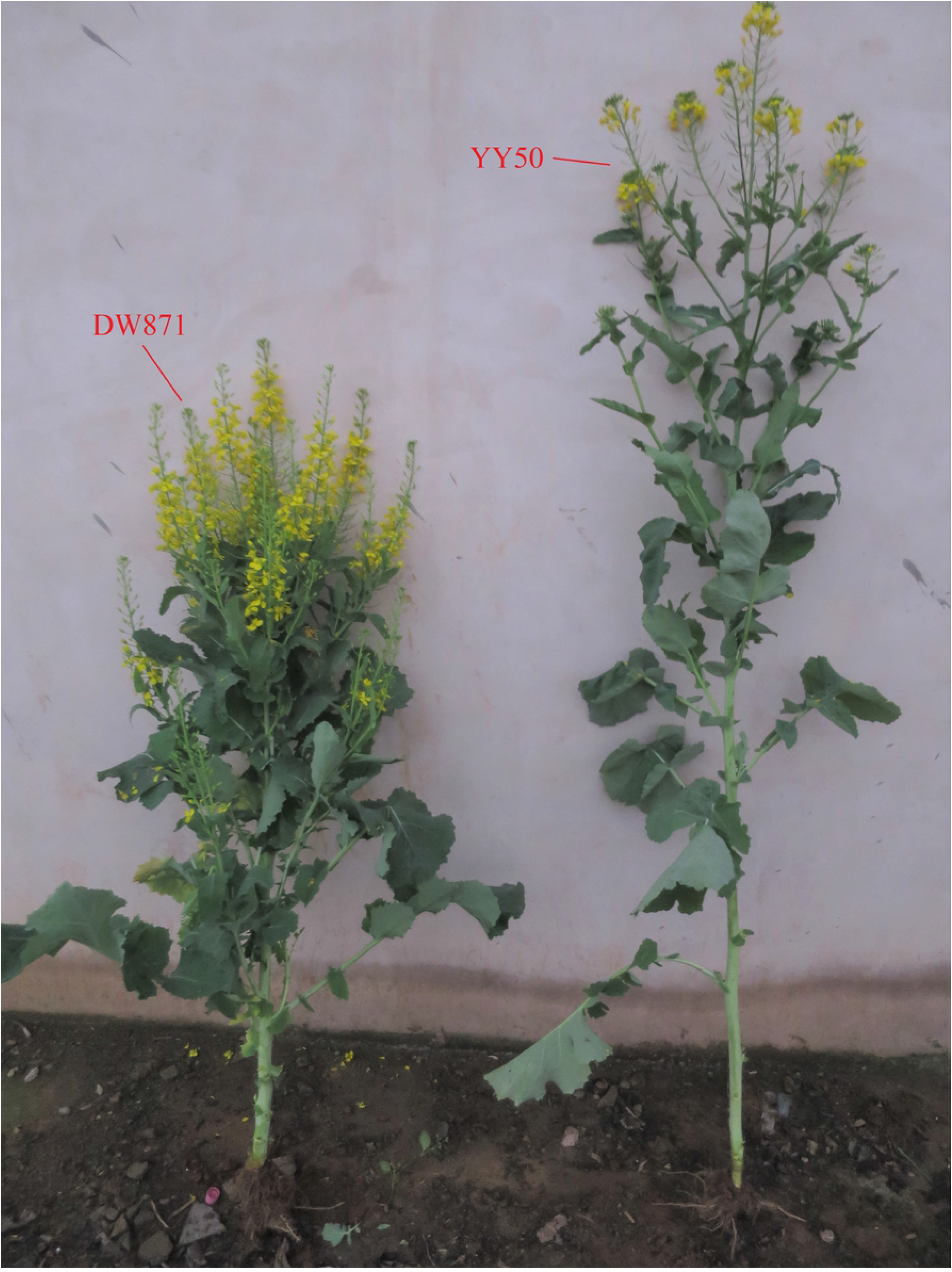
Comparison chart of DW871 and yy50

The method of transplanting seedlings is adopted environment that row spacing is 40cm, the plot is 20m^2^, and the density is 300,000 plants/hm^2^. The fertility level before and after transplanting is consistent with the field pest management.

### 2.2 Extraction of whole genome DNA and construction of plant height gene pool

According to the traits of field plants, the whole genome of 30 plants of high-stalk material and 30 plants of short-stalk material were extracted. After mixing according to the same ratio, gene pools for dwarf population (312bulk1) and a gene mixing pool for tall population (312bulk2) were constructed.

By the improved CTAB (Hexadecyl trimethyl ammonium Bromide) to extract the whole genome DNA. After quality and concentration identification, experiment adjust the DNA concentration, and finally mix with equal volume to establish two extreme plant height trait gene pools, respectively dwarf and tall gene traits^[35]^.

### 2.3 Construction of DNA library

Collect 30 leaves of high-stalk and short-stalked materials respectively to ensure that the leaves are intact and free of disease or damage. And quickly process plant materials at −80°C. And use the improved CTAB method to extract the whole genome DNA of the test material. Detect the quality of the whole genome DNA to ensure the next step. Build a PE library for each sample DNA. Then we randomly break the genomic DNA of each sample to 300-500bp fragments, connect the sequencing adapters, and prepare DNA clusters. Select the sequencing strategy PE150 on Illumina HiSeq X-ten (China, Illumina Company) to sequence the whole genome. In the end, a high-depth sequencing result of a mixed pool of single-strain DNA with extreme traits was obtained.

### 2.4 Compare clean data with reference sequence and detect variation

Obtaining the high-quality clean date helps to improve the quality of the data and the credibility of the follow-up results^[36]^. By performing data quality control on the library to obtain one clean date by Cutadapt (version 1.13) and Trimmomatic (version 0.36). Then compare the clean date to the reference genome (Darmor (v4.1)) by BWA (version 0.7.15-r1140), which is downloaded from www.genoscope.cns.fr/brassicanapus/data/)^[37]^. In order to obtain the BAM file for variant calling, we use the comparison result in SAM format to convert it through Samtoools and Picard tools. Finally, detecting mutations by GATK, we get candidate SNPs and InDels^[38]^.

### 2.5 Determine the candidate interval for dwarfing genes

In order to determine the candidate interval of dwarfing genes, we screen the genome based on Δ (SNP-index). And through the G’ value, we can effectively eliminate the “high frequency” noise associated with sequence variation. Get narrower candidate section. The formula^[25]^ for calculating the value of Δ(SNP-index) and G’ is (1)–(5):

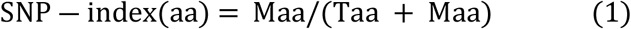

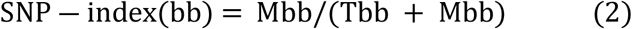

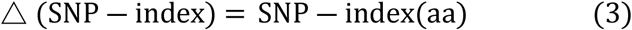

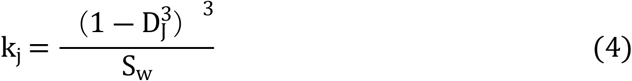

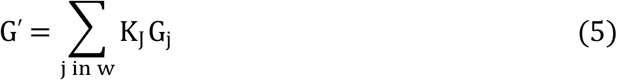

In the above formula, M represents the plant height trait, and T represents the highest marker, and aa and bb represent the genotypes of 312bulk1 and 312bulk2, respectively. Maa and Mbb represent the concentrations of the aa population derived from M and the bb population of M, and Taa and Tbb represent the depth of aa and T bb derived from T, respectively. Δ (SNP-index) indicates the degree of correlation between the marker and the trait. The closer Δ (SNP-index) is to 1, the stronger the correlation between the marker and the trait. D_j_ is the standard distance, S_w_ is the sum of (1-D_j_^3^)^3^ of all SNPs in W, G_j_ represents the average number of G mutants on related SNPs, and G’ is the weighted average of G_j_. In the actual operation process, we use QTLseqr (v0.7.3) to process the above formula and set the size of the sliding window to 2Mb. In addition, when analyzing the two sample pools according to the reading depth to verify the significance level of the relevant SNP, it needs to meet P<0.01.

In the process of determining the candidate interval, the conditions used are as follows: (1) The sequencing depth of the two parents is greater than 8; (2) The sequencing depth of the two pools is greater than 10; (3) The SNP-index value of the two pools cannot be greater than 0.8 or less than 0.2 at the same time;(4) Polymorphism between parents.

Based on the SNP detected by BSA, we use the principle of map-based cloning^[39]^. We will manually select 23 SNPs under the condition that the target range is covered as uniformly as possible and specific amplification primers can be designed. Capture the target DNA fragment by designing primers to establish a PE library. Secondly, based on the sequencing data of each site, we analyze the genotype of each target site, find the Individual exchanges among them, and determine the effective Individual exchange. Finally, comprehensively analyze multiple effective exchange individual plants to determine the candidate interval of the target gene.

## 3. Conclusion

### 3.1 Sequencing data and statistics of whole genome comparison results

This experiment was carried out in Hubei GENOSEQ Technology Co., Ltd., using Illumina Hiseq X-ten to sequence the DNA of two samples. In order to ensure the smooth progress of the follow-up work, remove the clean reads of low quality, containing joint boxes, and containing more N. A total of 192.80Mb sequencing data was obtained, 312bulk1 pool totaled 82.95Mb, and 312bulk2 pool totaled 109.85Mb. The average GC content of the two pools was 38.07%, and the average Q30 ratio was 92.71% (Table 1). The nucleobase quality of most sequencing sites in samples 312bulk1 and 312bulk2 are above Q30, and the nucleobase content is balanced, and the sequencing results are stable.

**Table 1:** Sample sequencing data

| Sample | Reads | Bases | GC(%) | Q20(%) | Q30(%) |
| --- | --- | --- | --- | --- | --- |
| 312bulk1 | 82,954,523 | 23,852,763,079 | 37.99 | 97.50 | 92.53 |
| 312bulk2 | 109,848,541 | 31,636,636,178 | 38.14 | 97.63 | 92.88 |
| Mean | 96,401,532 | 27,744,699,629 | 38.07 | 97.57 | 92.71 |
| Sum | 192,803,064 | 55,489,399,257 | - | - | - |

Process the sample DNA through the Illumina library to obtain PE reads (paired-end reads). Then the PE reads were compared with the *Brassica napus* L. reference genome (Darmor(v4.1)). The ratio of sample 312bulk1 to the reference genome sequencing data was 92.82%, the average coverage was 74.80%, and the average coverage depth was 17.14X; the comparison ratio of sample 312bulk2 was 93.34%, the average coverage was 73.88%, and the average coverage depth was 22.84X (Table 2). The contrast ratio of the sample is higher than 70%, and the coverage depth is greater than 10, which meets the requirements. The sequencing of the two samples has good randomness, and the sequencing data achieve uniform coverage of the reference genome.

**Table 2:**
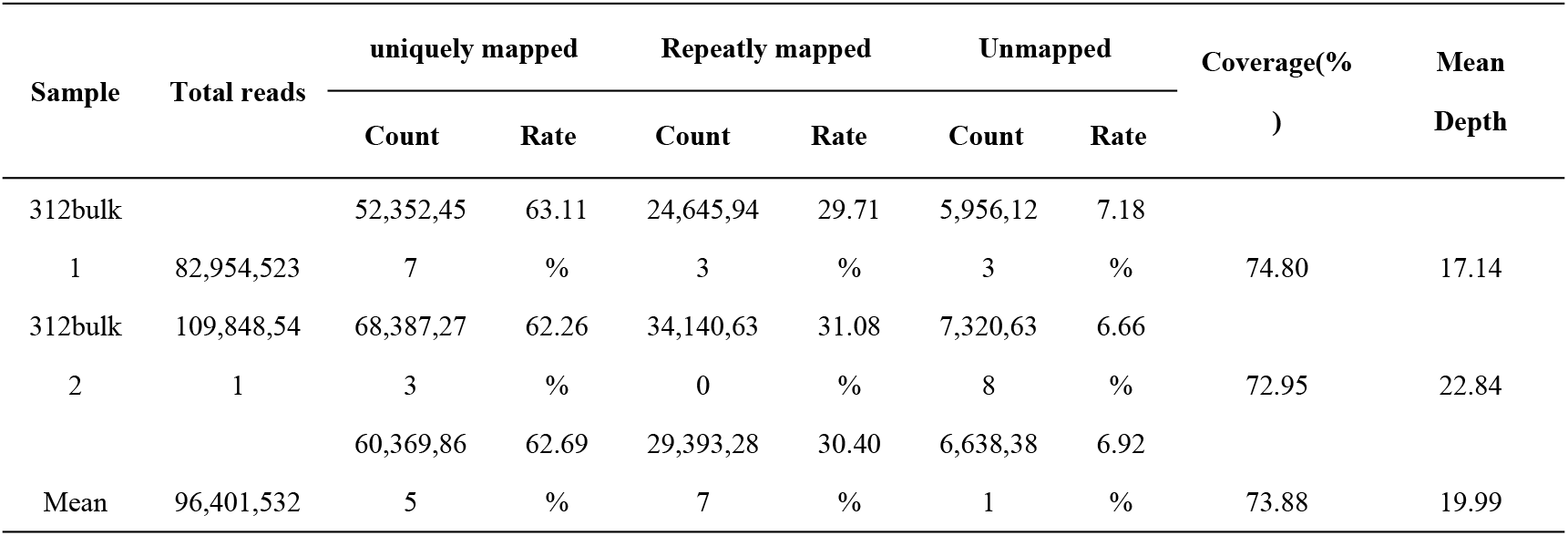
Statistics of comparison results of sequencing data

### 3.2 Screening of mutation sites by SNPs and InDels

Using GATK to complete the mutation detection of the sample DNA, we tested a total of 645.40Mb of data, and screened out 1,189,864 usable SNPs and 217,989 usable InDels. The labeling density of SNPs and InDels on A10 was 2847.13 and 682.83, and the labeling density was the highest in each chromosome (Table 3). Through the distribution map of available SNP/InDel sites on each chromosome, we can intuitively see that the markers are concentrated on A01 and A10 (Fig 2). It indicates that the markers associated with genes related to dwarfing traits of *Brassica napus* L. mainly exist on chromosome A10.

**Fig 2:**
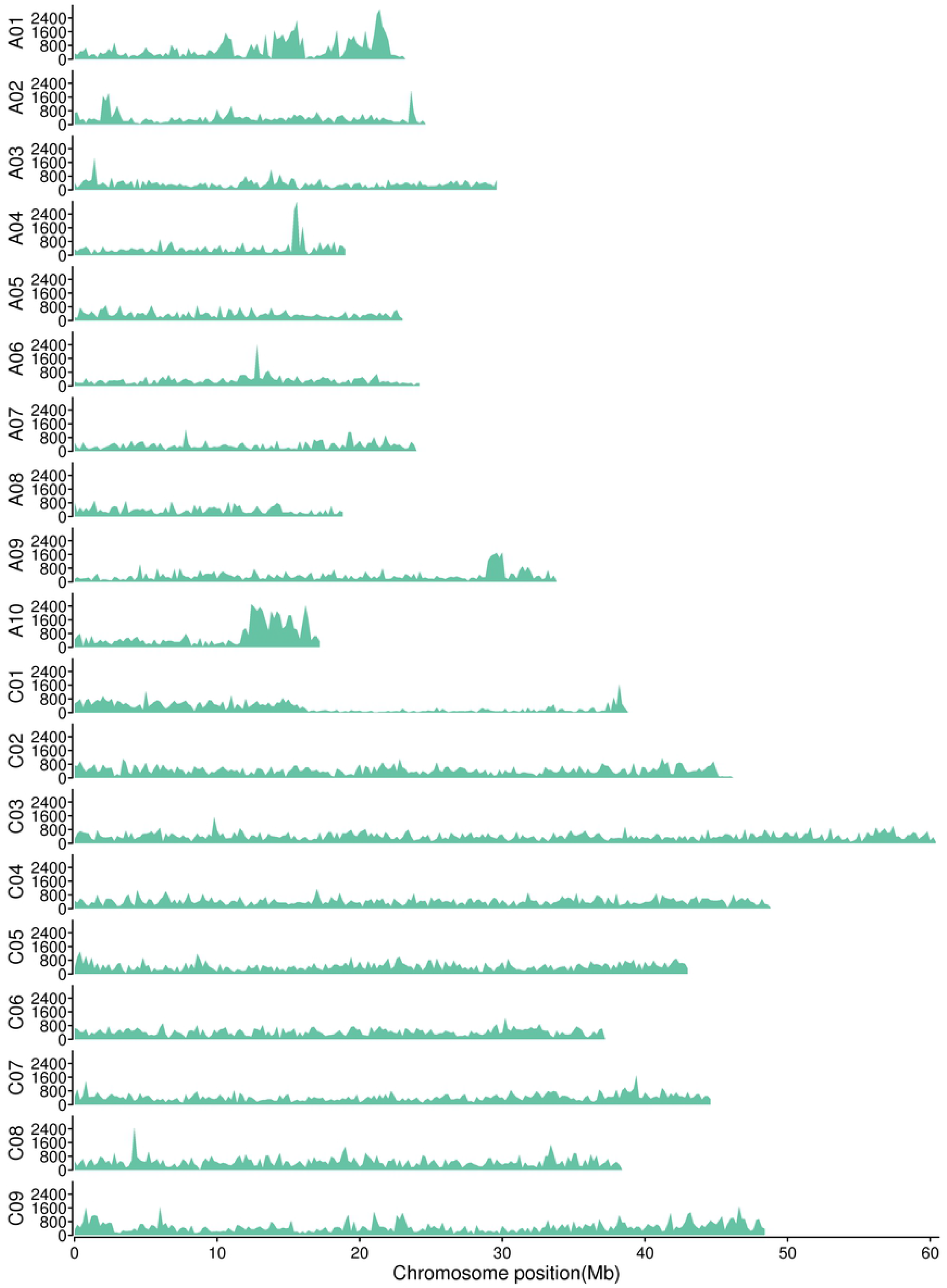
The x-axis is the chromosome position, and the y-axis is th

**Table 3:** Samples can be chromosome-divided statistics by SNP and InDel

| Chr | Length | SNPs |  | INDELs |  |
| --- | --- | --- | --- | --- | --- |
|  |  | Count | Density (Mb<sup>-1</sup>) | Count | Density (Mb<sup>-1</sup>) |
| A01 | 23,267,856 | 59,509 | 2557.56 | 14,075 | 604.91 |
| A02 | 24,793,737 | 39,798 | 1605.16 | 8,346 | 336.62 |
| A03 | 29,767,490 | 42,583 | 1430.52 | 10,389 | 349.00 |
| A04 | 19,151,660 | 32,691 | 1706.95 | 7,275 | 379.86 |
| A05 | 23,067,598 | 36,457 | 1580.44 | 7,209 | 312.52 |
| A06 | 24,396,386 | 34,581 | 1417.46 | 6,266 | 256.84 |
| A07 | 24,006,521 | 34,665 | 1443.98 | 7,478 | 311.50 |
| A08 | 18,961,941 | 30,037 | 1584.07 | 6,285 | 331.45 |
| A09 | 33,865,340 | 51,974 | 1534.73 | 10,964 | 323.75 |
| A10 | 17,398,227 | 49,535 | 2847.13 | 11,880 | 682.83 |
| C01 | 38,829,317 | 50,388 | 1297.68 | 9,425 | 242.73 |
| C02 | 46,221,804 | 92,105 | 1992.67 | 14,362 | 310.72 |
| C03 | 60,573,394 | 110,836 | 1829.78 | 19,723 | 325.61 |
| C04 | 48,930,237 | 94,559 | 1932.53 | 14,715 | 300.73 |
| C05 | 43,185,227 | 87,025 | 2015.16 | 13,040 | 301.96 |
| C06 | 37,225,952 | 72,051 | 1935.50 | 11,709 | 314.54 |
| C07 | 44,770,477 | 85,837 | 1917.27 | 13,645 | 304.78 |
| C08 | 38,477,087 | 81,256 | 2111.80 | 13,556 | 352.31 |
| C09 | 48,508,220 | 103,959 | 2143.12 | 17,647 | 363.79 |
| Whole | 645,398,471 | 1,189,846 | 1835.97 | 217,989 | 352.97 |

### 3.3 Preliminary positioning of candidate intervals

We determined the interval of Δ (SNP-index)>4 (Fig 3) and G’ value>2 (Fig 4), and determined the common interval between the two after comprehensive analysis. We located the candidate interval related to DW871 plant height traits between BnaA10g21330D and BnaA10g26440D on ChrA10. Starting at 14.70Mb and ending at 16.86Mb, the length of the candidate interval is 2.16Mb. In this range, we screened a total of 14,340 variants in the 312bulk1 and 312bulk2 genomes, including 512 genes, and 290 genes have large effects.

**Fig 3:**
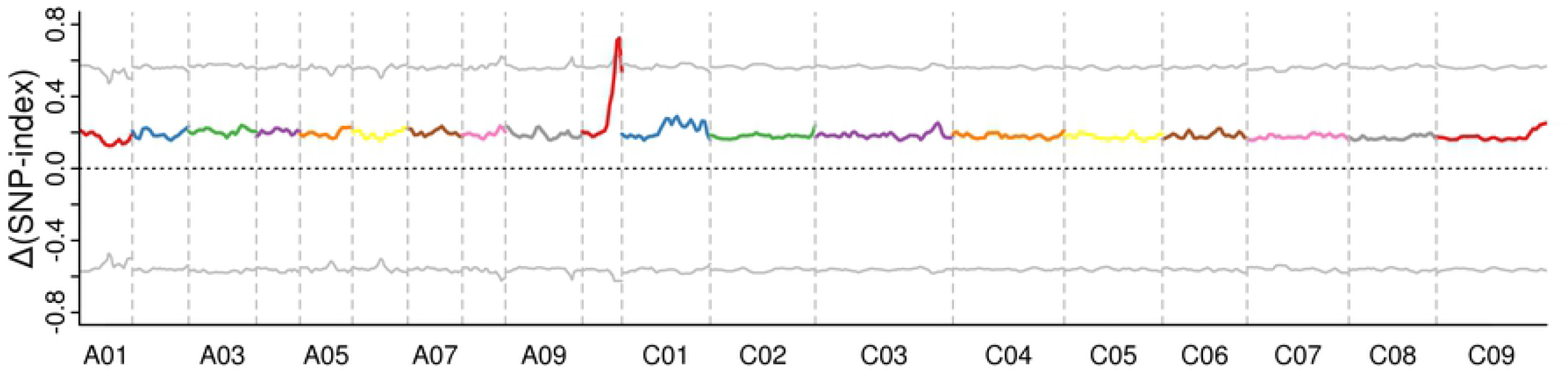
□ (SNP-index)

**Fig 4:**
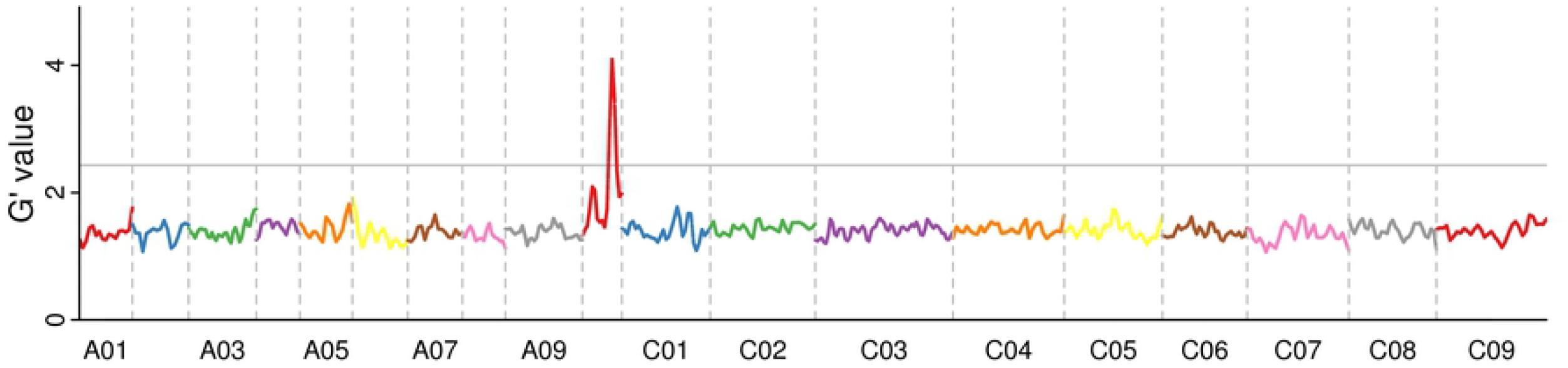
G’ value

### 3.4 Fine positioning of candidate intervals

The candidate intervals are between BnaA10g21330D and BnaA10g26440D on ChrA10. According to the method in 2.5, we sequenced a total of 500 samples and obtained a total of 13.47Mb clean PE reads. The GC value is 42.42, the Q30 value is 98.03, and the average coverage depth of the site is 2453.76X. Meet the requirements.

In order to further narrow the candidate interval, we need to determine the individual in the sample that has exchanged genotypes and phenotypes. Screening polymorphisms between dwarf and tall stalks, let us finally obtain 17 pairs of primers (Table 4, middle box). We analyzed the phenotypic composition of the samples and identified 13 effective exchange individual plants out of 500 samples.

**Table 4:**
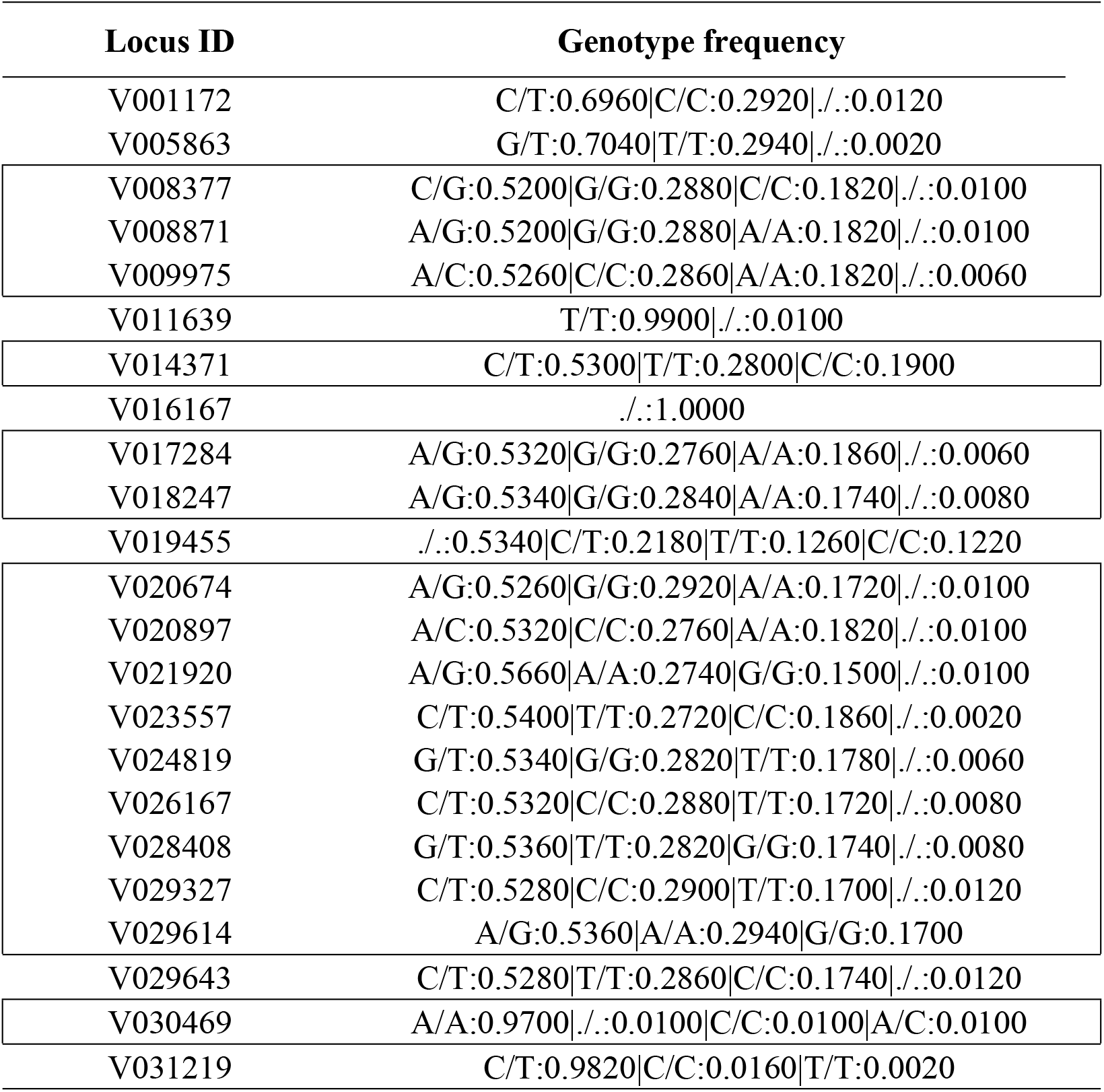
Statistics of genotype composition of target locus

| Locus ID | Genotype frequency |
| --- | --- |
| V001172 | C/T:0.6960\|C/C:0.2920\|./.:0.0120 |
| V005863 | G/T:0.7040\|T/T:0.2940\|./.:0.0020 |
| V008377 | C/G:0.5200\|G/G:0.2880\|C/C:0.1820\|./.:0.0100 |
| V008871 | A/G:0.5200\|G/G:0.2880\|A/A:0.1820\|./.:0.0100 |
| V009975 | A/C:0.5260\|C/C:0.2860\|A/A:0.1820\|./.:0.0060 |
| V011639 | T/T:0.9900\|./.:0.0100 |
| V014371 | C/T:0.5300\|T/T:0.2800\|C/C:0.1900 |
| V016167 | ./.:1.0000 |
| V017284 | A/G:0.5320\|G/G:0.2760\|A/A:0.1860\|./.:0.0060 |
| V018247 | A/G:0.5340\|G/G:0.2840\|A/A:0.1740\|./.:0.0080 |
| V019455 | ./.:0.5340\|C/T:0.2180\|T/T:0.1260\|C/C:0.1220 |
| V020674 | A/G:0.5260\|G/G:0.2920\|A/A:0.1720\|./.:0.0100 |
| V020897 | A/C:0.5320\|C/C:0.2760\|A/A:0.1820\|./.:0.0100 |
| V021920 | A/G:0.5660\|A/A:0.2740\|G/G:0.1500\|./.:0.0100 |
| V023557 | C/T:0.5400\|T/T:0.2720\|C/C:0.1860\|./.:0.0020 |
| V024819 | G/T:0.5340\|G/G:0.2820\|T/T:0.1780\|./.:0.0060 |
| V026167 | C/T:0.5320\|C/C:0.2880\|T/T:0.1720\|./.:0.0080 |
| V028408 | G/T:0.5360\|T/T:0.2820\|G/G:0.1740\|./.:0.0080 |
| V029327 | C/T:0.5280\|C/C:0.2900\|T/T:0.1700\|./.:0.0120 |
| V029614 | A/G:0.5360\|A/A:0.2940\|G/G:0.1700 |
| V029643 | C/T:0.5280\|T/T:0.2860\|C/C:0.1740\|./.:0.0120 |
| V030469 | A/A:0.9700\|./.:0.0100\|C/C:0.0100\|A/C:0.0100 |
| V031219 | C/T:0.9820\|C/C:0.0160\|T/T:0.0020 |

We verified the individual exchanges obtained after screening through these molecular markers, and found that the individual exchanges with high stalk phenotype decreased by 1, at the marker V008377, decreased 1, V008871 decreased 1, V009975 decreased 2, V014371 decreased 3, V017284 decreased 4; on the other side of the interval, the individual exchanges with the tall stalk phenotype are 1 at V029643 and V029614. This result shows that the target gene is between the markers V017284 and V029614. Comprehensive analysis of multiple individual exchange, we can determine the target gene between V018247 and V029614. Start at 15.51Mb and end at 16.60Mb. Its physical distance is 1.09Mb.

Between V018247 and V029614, we identified 231 genes. Through blasts in NCBI, 77 of which are genes, we are currently unable to determine their specific functions. In addition, we identified 24 genes related to transcription, 12 genes related to resistance, 42 genes involved in protein kinase recombination, 43 genes involved in intracellular metabolism, and 15 genes involved in cell growth and the apoptosis of process, and 5 genes related to protein modification. Some genes are related to Lignin anabolic pathway, petal growth and flowering factors.

## 4. Summary

The final purpose of a series of work by breeders is to obtain new varieties of rapeseed with high yield and mechanized harvesting. The continuous increase in the yield of rapeseed led to the appearance of lodging, which in turn caused the yield of rape to drop significantly, reducing the economic benefits of rape and the enthusiasm of farmers to plant. Therefore, the dwarf breeding of *Brassica napus* L. has become an important means to provide the mechanization rate of rapeseed production and increase farmers’ income. BSA-seq is a new generation method that relies on NGS to determine the candidate interval of the target gene. In this research, two pools with extreme differences in traits were constructed, named 312bulk1 and 312bulk2. The whole genome sequencing data of these two pools is all obtained by NGS sequencing after mixing multiple samples in equal proportions. After obtaining the sequencing data, by analyzing the data and calculating its SNP-index, Δ (SNP-index) and G’ value, we found the candidate interval of the DW871 dwarf gene on ChrA10 chromosome, 14.70Mb ~ 16.86Mb. The length of the candidate interval is 2.16Mb, including 512 genes, including 290 genes with large effects.

Because the current candidate interval is too large, it cannot meet the needs of subsequent gene mapping and breeding. Therefore, we need to further narrow the target gene candidate interval. SNP markers are a new generation of molecular markers and have been widely used. SNP genotyping technology is a gene mapping technology developed based on SNP molecular markers. Through map-based cloning and SNP genotyping technology, we can further narrow the candidate range of target genes. Through manual selection, we selected 23 SNP markers among the SNPs obtained after BSA, and finally obtained 17 polymorphic markers through polymorphism screening. Then through the comparison and analysis of the marker and the individual exchanges, we finally determined the candidate interval of the target gene, 15.51Mb ~ 16.60Mb, and its physical distance was 1.09Mb.

Between V018247 and V029614, we identified 231 genes. Among 77 genes, we are currently unable to determine their specific functions. In addition, we identified 24 genes related to transcription, 12 genes related to resistance, 42 genes involved in protein kinase recombination, 43 genes involved in intracellular metabolism, and 15 genes involved in cell growth and the apoptosis of process, and 5 genes related to protein modification. Some genes are involved in lignin anabolic pathway, petal growth and flowering regulation. Therefore, we conclude that the dwarfing trait may be related to the synthesis of lignin. In addition, because DW871 is a definite inflorescence, we speculate that the formation of dwarfing traits is related to the formation of definite inflorescence. This research provides a reference for further determining the target range of dwarfing genes and verifying the formation mechanism of related traits.

However, to determine the formation mechanism of dwarf traits and the cloning of dwarf genes still needs more in-depth work.

## Thanks

This research is co-sponsored by Guizhou science and technology key project(Qian Ke He Basis 〈2017〉 1415), Guizhou outstanding young science and technology talent project(Qian Ke He Platform Talent 〈2017〉 5642), Guizhou support plan project(Qian Ke He bracing 〈2017〉 2568), Guizhou RenCai JiDi(RCJD2018-14), Guizhou provincial engineering research center for plant conservation technology application, and Construction of laboratory platform of agricultural science and technology innovation center in karst mountains(Qian Ke Zhong Yin Di 〈2018〉4001). In addition, special thanks to Wen Dezhi of Guizhou Normal University for his selfless help in the translation and writing of this article.

